# The DEAD-box RNA helicase Dhx15 controls glycolysis and arbovirus replication in *Aedes aegypti* mosquito cells

**DOI:** 10.1101/2022.06.22.497281

**Authors:** Samara Rosendo Machado, Jieqiong Qu, Werner J.H. Koopman, Pascal Miesen

## Abstract

*Aedes aegypti* mosquitoes are responsible for the transmission of arthropod-borne (arbo)viruses including dengue and chikungunya virus (CHIKV), but in contrast to human hosts, arbovirus infected mosquitoes are able to efficiently control virus replication to sub-pathological levels. Yet, our knowledge about the molecular interactions of arboviruses with their mosquito hosts is largely incomplete. Here, we aimed to identify and characterize novel host genes that control arbovirus replication in *Aedes* mosquitoes. RNA binding proteins (RBPs) are well known to regulate immune signaling pathways in all kingdoms of life. We therefore performed a knockdown screen targeting 461 genes encoding predicted RBPs in *Aedes aegypti* Aag2 cells and identified 15 genes with antiviral activity against a Sindbis reporter virus. Amongst these, three DEAD-box RNA helicases, AAEL004419/Dhx15, AAEL008728 and AAEL004859 also acted as antiviral factors in dengue and CHIKV infections. Here, we explore the mechanism of Dhx15 in regulating an antiviral transcriptional response in mosquitoes by silencing *Dhx15* in Aag2 cells followed by deep-sequencing of poly-A enriched RNAs. *Dhx15* knockdown in uninfected or CHIKV-infected cells resulted in differential expression of 856 and 372 genes, respectively. Interestingly, amongst the consistently downregulated genes, *glycolytic process* was the most strongly enriched GO term as the expression of all core enzymes of the glycolytic pathway was reduced, suggesting that Dhx15 regulates glycolytic function. A decrease in lactate production supported the observation that *Dhx15* silencing functionally impaired glycolysis. Modified rates of glycolytic metabolism have been implicated in controlling the replication of several classes of viruses and strikingly, infection of Aag2 cells with CHIKV by itself also resulted in the decrease of several glycolysis genes. Our data suggests that Dhx15 regulates replication of CHIKV, and possibly other arboviruses, by controlling glycolysis in mosquito cells.

## Introduction

The yellow fever mosquito *Aedes aegypti* is the principal vector of medically important arthropod-borne viruses (arboviruses) such as Chikungunya virus (CHIKV; genus *Alphavirus*, family *Togaviridae*) and dengue virus (DENV; genus *Flavivirus*, family *Flaviviridae*) (1–3). CHIKV and DENV infections cause similar, flu-like symptoms including headache, fever and muscle pain. More serious CHIKV infections manifest with severe joint pain and arthritis that sometimes persist for weeks up to years (4, 5), whereas serious DENV infections may result in loss of body fluid and hemorrhagic fever (1). *Ae. aegypti* mosquitoes were originally restricted to (sub)tropical countries. However, elevated global temperatures, increased urbanization and more extensive international travel and trade have favored mosquito invasion of more temperate climate zones (1). The expansion of the *Ae. aegypti* habitat has consequently lead to the global spread of arboviruses alike (6).

The ability of mosquitoes to acquire, replicate and transmit arboviruses, collectively referred as vector competence, is a key determinant for efficient arbovirus transmission (7). Upon acquisition in an infected bloodmeal, viruses initially infect midgut epithelial cells and subsequently disseminate to secondary tissues. Once a systemic infection is established and high viral titers are reached in the mosquito saliva, arbovirus transmission takes place (7–9). Interestingly, virus accumulation in mosquitoes generally remains sub-pathological (10), suggesting that mosquitoes are able to efficiently reduce virus replication (resistance) and/or prevent virus-induced tissue damage (tolerance) (11). However, to date, a comprehensive picture of the molecular processes that control arbovirus replication in the mosquito hosts is still lacking (9).

The fruit fly *Drosophila melanogaster*, a well-established genetic model organism, has been instrumental in dissecting the genetic basis of antiviral immunity in insects (12–14). In *Drosophila*, the RNA interference (RNAi) pathway has been established as an important antiviral immune pathway that restricts both RNA and DNA viruses (15, 16). Studies in mosquitoes have confirmed the broad antiviral activity of this pathways across dipteran insects (17). Moreover, work in *Drosophila* has indicated that transcriptional responses through inducible immune signaling pathways contribute to antiviral immunity, in particular the JAK-STAT (Janus kinase-signal transducers and activators of transcription) pathway and the two NFκB (Nuclear factor κB)-related Toll and IMD (immune deficiency) pathways (18–20). Whereas RNAi destroys viral RNA directly, transcriptional regulation of immune responses has been proposed to up-regulate anti-microbial peptides (21) or module metabolic responses (22), but in general, the role of transcriptional responses in antiviral immunity in *Ae. aegypti* mosquitoes is still largely understudied (17).

Here, we set out to identify new genetic determinants that control mosquito immune responses focusing on RNA binding proteins (RBPs), which regulate signaling pathways in response to infection in all kingdoms of life (23–27). In particular, DEAD-box RNA helicases, a subgroup of RBPs (28), comprise well-known examples of enzymes that recognize viral RNA and modulate antiviral signaling (23–25). These include the cytoplasmic viral RNA sensors RIG-I (retinoic-acid-inducible gene I) and MDA5 (melanoma-differentiation-associated gene 5), which are key activators of interferon signaling in vertebrates (24), the antiviral RNAi effector Dicer-2 (29), and many other RNA helicases that act as co-receptors and signaling intermediates in diverse immune pathways (26, 27). Due to the important and versatile role of RBPs, we deemed it likely that members of this family control arbovirus replication in vector mosquitoes.

To identify RBPs that interfere with arboviruses replication in mosquitoes we performed a knockdown screen in *Ae. aegypti* Aag2 cells and assessed virus replication of a Sindbis reporter virus (SINV; genus *Alphavirus*, family *Togaviridae*). This approach uncovered fifteen antiviral genes that upon knockdown enhanced virus replication; amongst these, three DEAD-box RNA helicases, AAEL004419, AAEL008728 and AAEL004859 had broad antiviral activity against SINV, CHIKV and DENV.

We further characterized the mechanism underlying antiviral activity of AAEL004419, the mosquito orthologue of Dhx15. Knockdown of this helicase decreased the expression of genes involved in glycolysis and consequentially reduced lactate production in mosquito cells. Glycolysis is a key process in energy metabolism by converting glucose into pyruvate, which is taken up by the mitochondria, oxidized to acetyl-CoA, and further metabolized in the tricarboxylic acid (TCA) cycle. Under anaerobic conditions, pyruvate can be converted into lactate, which is released from the cell (30). Besides energy production, glycolysis provides the precursors for essential biomolecules including nucleotides, amino acids and glycolipids/proteins (30, 31). The activity of glycolysis has direct effect on antiviral responses and has been reported to change upon infection with distinct viruses (32, 33). In line with this notion, we show that CHIKV infection of Aag2 cells reduced the expression of several glycolysis related genes, similar to knockdown of AAEL004419/Dhx15. This crosstalk at the level of glycolytic gene expression suggests that AAEL004419/Dhx15 controls CHIKV infection by regulating the glycolysis pathway in mosquito cells.

## Materials and methods

### RNA binding proteins selection

Genes encoding RNA binding proteins were selected based on gene annotations from VectorBase release 2017-8 that used the *Ae. aegypti* L3 genome as reference genome. Using the Biomart-plugin, genes associated with the gene ontology (GO) term “RNA binding” (GO:0003723) were selected from the *Ae. aegypti* gene dataset. This analysis was repeated for four additional dipteran species with annotated genomes: *Ae. albopictus, Culex quinquefasciatus, Anopheles gambiae* and *Drosophila melanogaster*. For the predicted RNA binding proteins from these species, *Ae. aegypti* orthologues were identified using the Biomart functionality within VectorBase and all list of genes were combined into a non-redundant set of genes encoding putative RNA binding proteins. We manually excluded genes that were unambiguously annotated as part of the core transcriptional, translation and splicing machinery. The remaining genes were included in the RNAi screen and selected for double-stranded RNA production and knockdown in Aag2 cells (Table S1).

Of note, retrospective manual inspection of the candidate genes included in the screen identified a few genes not to contain canonical RBP domains. This may be due to the revisited genome annotation or the orthologue-conversion step which may define an *Ae. Aegypti* orthologue that lacks RBP domains. Also, due to several updates of the *Ae. aegypti* reference genome annotation, some genes initially selected have been discontinued from the database or the annotation has been changed. Throughout the manuscript, the current gene identifiers of the L5 version of the *Ae. aegypti* genome are used. *NB*: The Biomart-function within VectorBase has been discontinued and replaced with a different search interface.

### Cells

*Aedes aegypti* Aag2 cells and the C3PC12 clone derived from these cells (cleared of the persistently infecting viruses Cell fusing agent virus, Phasi Charoen like virus and Culex Y virus) were maintained at 28 °C in Leibovitz’s L-15 medium (Invitrogen: catalogue number: 21083027) supplemented with 10% foetal bovine serum (Gibco), 50 U/mL penicillin, 50 μg/mL streptomycin (Gibco), 2% tryptose phophate broth (Sigma), and 1% non-essential amino acids (Gibco). For lactate assays, Aag2 C3PC12 cells were cultured in Schneider’s *Drosophila* medium (Invitrogen, catalogue number 21720024) containing 11.11 mM D-glucose and 12.32 mM L-glutamine. This medium was supplemented with 10% foetal bovine serum (Gibco), 50 U/mL penicillin, and 50 μg/mL streptomycin (Gibco). Hela cells, BHK15 and BHK21 cells were maintained at 37 °C, 5% CO_2_ in Dulbecco’s modified Eagle medium (DMEM) (Life Technologies, catalogue number 11995065) containing 25 mM D-glucose, 4 mM L-glutamine, and 1 mM sodium pyruvate. This medium was supplemented with 10% foetal bovine serum (Gibco), 50 U/mL penicillin, and 50 μg/mL streptomycin (Gibco).

SINV-nLuc, expressing a Nano-luciferase (nLuc) reporter as fusion protein with the SINV non-structural protein 3 (nsP3), was prepared on BHK-21 cells as previously described (34). The CHIKV expression plasmid encoding the Leiden synthetic (LS3) wildtype strain (35) was kindly provided by Dr. M.J. van Hemert (Leiden University Medical Center) and viral RNA was obtained by *in vitro* transcription on linearized plasmids using T7 mMessage mMachine (Invitrogen). RNA was then transfected into BHK-21 to grow infectious virus. Stocks of DENV serotype 2 (New Guinea C [NGC] strain) were prepared on *Aedes albopictus* (C6/36) cells. For quantification of viral stocks, SINV and CHIKV were titrated on BHK-21 cells, and DENV2 was titrated on BHK-15 cells.

To determine infectious DENV titres upon helicase silencing, end-point dilution assays were performed. A day prior to the titration, 1×10^4^ BHK-15 cells were seeded per well in a 96-well flat bottom plate. For the titration, a 10-fold serial dilution of virus samples were added to the cells in quadruplicate. After an incubation time of 7 days, cells were inspected for cytopathic effect (CPE). The virus titre was calculated according to the Reed and Muench method (36).

### *Aedes aegypti* mosquito rearing and dissection

*Aedes aegypti* mosquitoes (Black Eye Liverpool strain, obtained from BEI resources) used for dissection were reared at 28 °C and 70% humidity with automated room lighting set at a 12:12 hours light/dark cycle. Larvae were fed with Tetramin Baby fish food (Tetra). Adult mosquitoes were fed with a 10% sucrose solution. Five days old female mosquitoes (n=30) were dissected as previously described (37). Entire mosquitoes or dissected tissues (ovaries, midgut, head, thorax, rest of the body) were homogenized in 300 μl RNA-Solv reagent (Omega Bio-Tek) using a Precellys 24 homogenizer (Bertin technologies). To the homogenates, 700 μl RNA-Solv reagent was added and total RNA was isolated according to manufacturer’s recommendation.

### Expression construct cloning

cDNAs of AAEL004859, Dhx15 and AAEL008728, were cloned into pUbGw and pU3Fw for N-terminal tagging with GFP or 3xFlag, respectively. The vector pUbGw was modified from the expression vector pUbB-GW, (kindly provided by Dr. ir. Gorben Pijlman, University of Wageningen), as previously described (38). The expression vector pU3Fw was derived from the pUbGw vector by exchanging the GFP sequence with a 3xflag tag (39). For AAEL004859 and AAEL008728, gene-specific primers were used to amplify the genes from Aag2 cDNA and insert these sequences into an intermediate cloning vector using the TOPO-TA cloning kit (Thermo Fisher) according to the manufacturer’s protocol. The obtained plasmids were used as template in a subsequent PCR for In-Fusion HD Cloning (Takarabio). The purified PCR products were inserted into the Gateway entry vector pDonor/Zeo vector (Invitrogen) using the In-fusion reaction according to the manufacturer’s protocol. The sequence of the entry vector was confirmed by Sanger sequencing and LR-recombination (Thermo Fisher) was performed to recombine the sequence of the genes of interest to the destination vectors pUbGw and pU3Fw. For Dhx15, PCR amplification with Gateway cloning compatible primers was performed directly on Aag2 gDNA using CloneAmp Hifi PCR pre-mix (Takara), without prior amplification in a TOPO TA cloning vector. The PCR product was inserted in the pDonor/Zeo entry vector and recombined into the destination vectors using the Gateway cloning protocol (Thermo Fisher) as described above. Primer sequences are provided in Table S1.

### dsRNA production

dsRNA targeting each of the 461 RNA-binding proteins or Argonaute-2 (Ago-2) and firefly luciferase as positive and negative control, respectively, were produced from T7 promoter flanked PCR products. The T7 sequence was either directly present in the primer sequence used to generate the PCR products or they were introduced during a second PCR step using T7 universal primers that hybridize to short GC-rich tags that were introduced to the PCR products in the first PCR (see Table S1 for primer sequences). These PCR products were *in vitro* transcribed using a homemade T7 polymerase enzyme. For the formation of double-stranded RNA, the reactions were heated to 90 °C for 10 minutes and then allowed to gradually cool to room temperature. To purify the dsRNA, GenElute Mammalian Total RNA kit (Sigma) or GenElute 96 Well Total RNA purification Kit (Sigma) was used according to the manufacture’s protocol.

### Transfection of dsRNA and infection of Aag2 cells

For silencing experiments, Aag2 cells were seeded at a density of 1.5×10^5^ cells/well in a 24-wells plate or 5×10^4^ cells/well in a 96-wells flat bottom opaque white plate. For each condition, 3 wells were seeded 24 hrs prior to the first dsRNA transfection. In the 24-wells plate format, transfection mixes containing 300 μl non-supplemented L-15 medium, 450 ng dsRNA and 1.8 μl X-treme GENE HP DNA transfection reagent (Sigma) were prepared according to the manufacturer’s instructions. Per well, 100 μl of the transfection mix was added in a dropwise manner. For the 96-wells plate format, the volumes and amounts of the components of the transfection mix was one third of the quantities used for 24-wells plates. Three hours post-transfection, the medium was replaced with supplemented L-15 medium. To enhance knockdown efficiency, transfection was repeated 48 hours after the first transfection.

Where indicated, Aag2 cells were virus infected at the indicated multiplicity of infection (MOI) when changing the medium after the second transfection and cells were harvested 48 hours post-infection for downstream analyses.

### Cell fractionation

For plasmid transfection experiments, Aag2 cells were seeded 24 hrs prior to transfection at a density of 3.7×10^6^ cells/well in a 6-well plate. For each reaction, transfection mixes were prepared containing 500 μl non-supplemented L-15 medium, 5 µg plasmid DNA (Flag-tagged helicases) and 5 μl X-treme GENE HP DNA transfection reagent. Where indicated cells were infected with SINV after the transfection and samples were harvested 48h post infection. For sample preparation, Aag2 cells were resuspended, washed with PBS and pelleted at 300 x *g* for 5min. Next, cell pellets were lysed using cytoplasmic lysis buffer (50 mM NaCl, 25 mM Tris-HCl pH 7.5, 2 mM EDTA, 1x protease inhibitor, 0.5% NP40) and the cytoplasmic and the nuclear fractions were separated after 10 minutes centrifugation at 9600 x *g* at 4 °C. To the supernatant (cytoplasmic fraction) 5x Laemmli buffer (4% SDS, 0.004% bromophenol blue, 0.125 M Tris-HCl pH 6.8, 20% glycerol, 10% 2-mercaptoethanol) was added to a final concentration of 1x, the nuclear pellet was resuspended in Laemmli buffer diluted to 1x in cytoplasmic lysis buffer. For western blot, lysate fractions representing equal number of cells were loaded on gel.

### Co-immunoprecipitation

For co-transfection, 2.2×10^7^ Aag2 cells were seeded in a T-75 flask. To the transfection reaction for co-immunoprecipitation 30 µl of each plasmid DNA (GFP- and Flag-tagged helicases) and 60 µl X-treme GENE HP DNA transfection reagent was added. After two and a half hours incubation at 28 °C, the medium containing transfection reagents was replaced with supplemented L-15 medium.

Aag2 cells co-expressing GFP- and Flag-tagged RNA helicases were lysed in RIPA buffer (1% Triton X-100, 150 mM NaCl, 0.1% SDS, 0.5% Na-deoxychelate, 50 mM Tris pH 8.0, 1x protease inhibitor). The lysate was subjected to affinity enrichment using magnetic GFP-TRAP beads (ChromoTek) following the manufacture’s protocol. Beads were washed in washing buffer (10 mM Tris/Cl pH 7.5, 150 mM NaCl, 0.5 mM EDTA, 1x complete-EDTA free, and 1 mM PMSF). Where indicated, the samples underwent RNase A (Thermo Fisher) treatment for 7.5 minutes at 37 °C. After RNase A treatment, at least one additional washing step preceded the final elution. To the input samples taken before the precipitation, samples of washing steps, and the final eluate 5x Laemmli buffer diluted to 2x was added. Samples were heated at 90 °C for 10 minutes and analysed using western blot.

### Western blotting

For western blotting, protein samples were separated on polyacrylamide gels, blotted to nitrocellulose membranes and probed with the indicated antibodies. The primary antibodies used were mouse anti-H3K9me2 (Abcam ab1220), rat anti-α-tubulin (Bio-Rad), mouse anti-Flag M2 (Sigma), and rat anti-GFP (ChromoTek). The secondary antibodies used were: IRdye680 or IRdye800 conjugated goat anti-rat or goat anti-mouse (LI-COR). Primary antibodies were diluted 1:1000, and secondary antibodies 1:10000. Western blots were imaged on an Odyssey CLX imaging system (LI-COR).

### RNA isolation

Aag2 cells were homogenized in RNA-Solv reagent (Omega Bio-Tek) and RNA extraction was performed as described in the manufacturer’s instructions. Briefly, to 1 mL RNA-Solv reagent, 200 μl of chloroform was added and thoroughly mixed. After centrifugation, the aqueous phase was collected, and RNA was precipitated using isopropanol. This mix incubated for 1 hour at 4 °C followed by centrifugation to pellet the RNA. Pellets were washed twice in 80% ethanol, dissolved in nuclease free water, and quantified using a Nanodrop spectrophotometer.

### Reverse transcription and (quantitative) PCR

For reverse transcription followed by quantitative polymerase chain reaction (RT-qPCR), 1 μg of RNA was DNase I (Ambition) treated according to the manufacturer’s protocol and reverse transcribed using the TaqMan MultiScribe Reverse Transcription Kit (Applied Biosystems) using poly-dT and random hexamer primers. Quantitative PCR was performed on a LightCycler 480 (Roche) using GoTaq qPCR Mix (Promega), according to the manufacturer’s protocol. Relative expression of target genes were calculated using the 2^(-ΔΔCT)^ method (40) for which the expression of lysosomal aspartic protease (LAP) was used as an internal reference. End-point PCR to detect gene expression in mosquito tissues was performed using GoTaq polymerase (Promega) according to the manufacturer’s instructions. Sequences of primers are indicated in Table S1.

### Luminescence and Cell viability assay

Renilla-Glo Luciferase assay (Promega) was used to quantify nLuc reporter activity. The recommended volumes indicated in the manufacturer’s protocol was adapted and 70 µl of the reconstituted Renilla-Glo luciferase reagent was used per well of the 96-well plate. The CellTiter-Glo 2.0 assay (Promega) was used to quantify viable cells, according to the manufacturer’s instructions. For both assays, luminescence was measured on a Perkin Elmer Counter Victor 3 plate reader.

### RNA-sequencing library preparation and analysis

TruSeq Stranded mRNA kit (Illumina) was used for library preparation from total RNA according to the manufacturer’s protocol. The input for the library preparation was 1 µg RNA to obtain double-stranded cDNA. The prepared libraries were quantified and controlled for sample quality using a DNA1000 Bioanalyzer (Agilent). Next, the libraries were sequenced using Illumina HiSeq 4000 sequencing (GenomEast Platform).

### Bioinformatics analyses

After initial quality control by the sequencing platform, raw sequence reads were aligned to the *Aedes aegypti* LVP_AGWG AaegL5.1 reference genome (retrieved from VectorBase) using STAR 2.5.0 (41) with default settings. Detailed summary of the RNA-seq data can be found in Table S3. R package DESeq2 (42) using read count per gene was used for statistical analysis of differential gene expression (with adjusted *P* value < 0.05) and principal-component analysis. Genes were considered expressed if the mean of the DESeq2-normalized counts (baseMean) was higher than 10. The R package pheatmap (RRID:SCR_016418) was used to generate the heatmap for differentially expressed genes upon CHIKV infection, which was based on z-scores of normalized gene expressions (log10FPKM). The heat maps showing differential expression of glycolytic genes (based on log2-transformed fold changes) were generated in Microsoft Excel using three colour scale option of the conditional formatting function. Expression analysis of helicases in published datasets was performed as described previously (43, 44). Briefly, publicly available datasets were retrieved from NCBI Sequence Read Archive and mapped to the AaegL5 genome using STAR aligner version 2.5.2b (41). Raw read counts were then normalized with DESeq2 (42) and plotted with ggplot2 (45). GO term enrichment analysis was performed using DAVID (Database for Annotation, Visualization and Integrated Discovery) (46, 47). The STRING database was used to predict protein-protein interactions (48). Domain structure of RNA helicases was retrieved from Simple Modular Architecture Research Tool (SMART) (http://smart.embl-heidelberg.de/).

Phylogenetic analysis of RNA helicases was performed using the Multiple sequence alignment tool available on GenomeNet operated by the Kyoto University Bioinformatics Center (https://www.genome.jp/tools-bin/clustalw). As input, the protein sequences of the DEAD domains of *D. melanogaster* and *Ae. aegypti* DEAD-box RNA helicases were used. DEAD box helicases were identified using the “Search for” function in VectorBase asking gene identifiers based on InterPro Domain database. The specific domain to be searched for was set to PF00270: DEAD DEAD/DEAH box helicase domain. The resulting gene lists were obtained for both *D. melanogaster* and *Ae. aegypti* and the ‘edit-orthologues’ function was used to identify orthologous genes in the other species, respectively. The obtained lists were compiled into one non-redundant gene list of DEAD-box helicases for each species. Amino acid sequences of the DEAD domain of each protein were retrieved from the SMART database, or if unavailable, manually extracted from the protein sequences using the amino acid coordinates given by PFAM. The maximum likelihood tree was generated on the multiple sequence alignment using the FastTree full algorithm in GenomeNet, which is based on FastTree 2 (49).

### 2-deoxy-D-glucose treatment and lactate concentration measurement

Aag2 cells cultured in Schneider’s *Drosophila* medium and Hela cells cultured in DMEM medium were seeded 24 hours prior to 2-deoxy-D-glucose (Sigma) treatment. Cells were incubated for 48 hours with either 24 mM or 50 mM 2-deoxy-D-glucose, harvested, and samples were analysed using a lactate assay kit (Sigma-Aldrich). The concentration used for 2-deoxy-D-glucose was experimentally optimized in house. Lactate concentration was measured using a colorimetric detection following the manufacturer’s instructions.

### Statistical analysis

Unless indicated differently, experiments had three biological replicates and the data are represented as mean +/- standard deviation. Statistical significancy was attributed when *p*-value was <0.05. Graphs and statistical analysis were generated using GraphPad Prism (version 8.0.0 for Windows).

## Results

### A targeted RNAi screen in mosquito cells identifies novel host genes that control SINV replication

To identify RBPs that control virus replication in *Ae. aegypti*, we designed a targeted knockdown screen in Aag2 cells. Using the biomart plugin in VectorBase (release 2017-8), we selected all genes from the *Aedes aegyp*ti L3.3 genome annotation that were associated with the GO term RNA binding (Accession GO:0003723). We also identified *Ae. aegypti* orthologues of predicted RBPs in other mosquito species (*Ae. albopictus, Culex quinquefasciatus* and *Anopheles gambiae*) as well as the fruit fly *Drosophila melanogaster* and combined all datasets into a non-redundant list of 635 genes. We manually excluded 132 genes that were part of the core transcription, splicing and translation machineries. Another 42 genes were omitted because the PCR amplification to generate the template for *in vitro* transcription repeatedly failed. Overall, we managed to successfully produce double-stranded RNA (dsRNA) for knockdown of 461 genes, which represent the set of genes included in the first screening round (Table S2).

All genes were individually silenced in Aag2 cells followed by infection with a recombinant Sindbis virus expressing a nano-luciferase reporter gene as a fusion protein with nsP3 (34) (Fig. 1A). In the initial screening round, knockdown of 38 and 49 genes resulted in an ≥ 2-fold increase or decrease of luciferase levels, respectively, compared to the non-targeting control knockdown (Fig. 1B and C). We repeated the knockdown experiment for these genes using the same dsRNA preparation and, for those that were reproducible, we generated a second set of dsRNA targeting a different region of each gene to account for possible off-target effects (Fig. 1D, Table S2). As controls, we included silencing of the antiviral RNAi core factor Ago2 and knockdown of the SINV genomic RNA itself. With this extensive confirmation procedure, we validated the phenotype of fifteen antiviral hits (Fig. 1D) and four proviral hits (Table S2).

**Figure 1:**
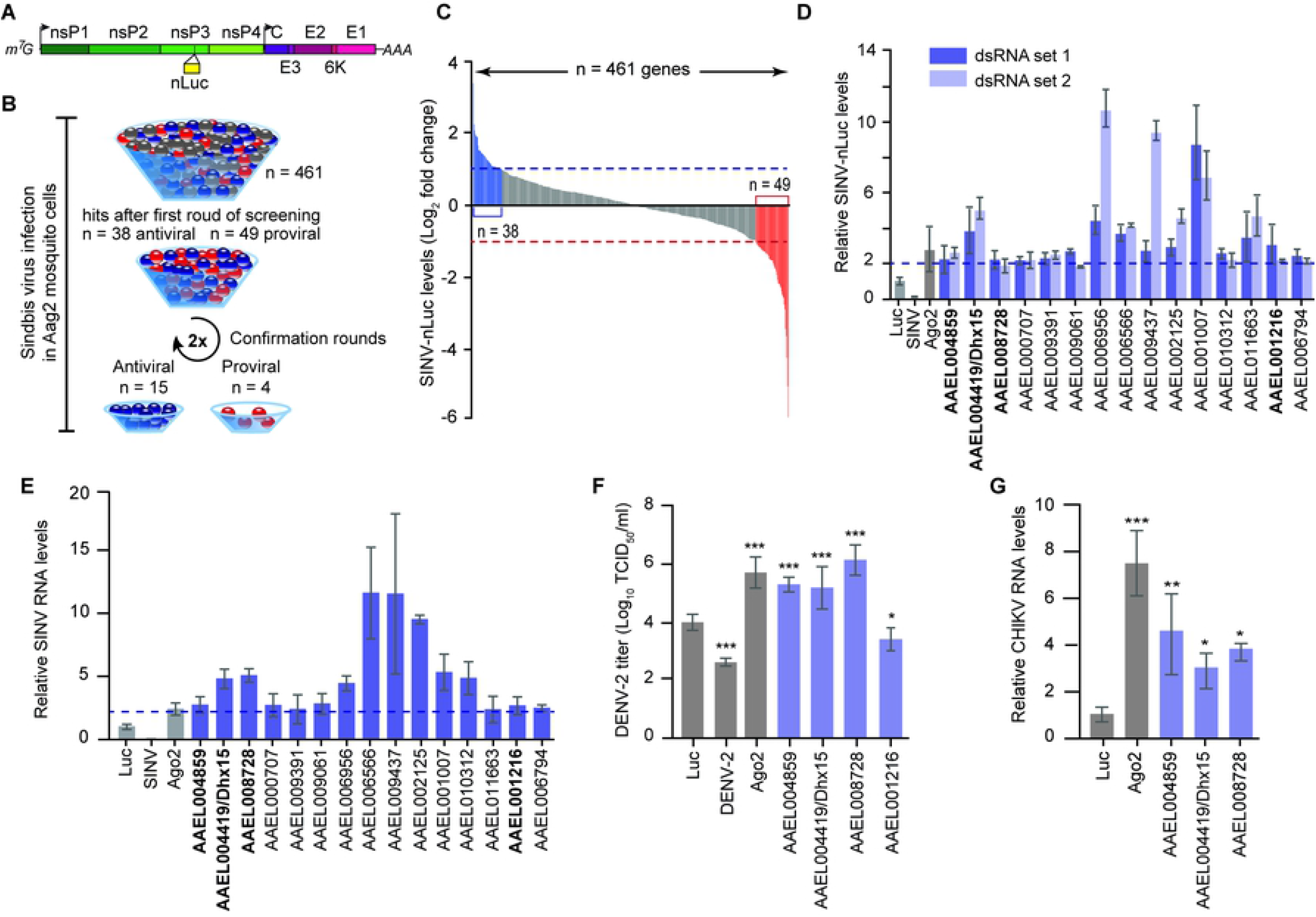
RNAi screen identifies RNA-binding proteins (RBPs) that control arboviruses replication in mosquito cells. **A)** Schematic representation of recombinant Sindbis virus expressing a nano-luciferase reporter gene as a fusion protein with nsP3. The individual non-structural and structural viral proteins are depicted in different shades of green and purple, respectively. The position of the nLuc is marked by the yellow bar. **B)** Schematic flow of the RNAi screen. Antiviral, proviral and neutral genes were depicted in blue, red and gray respectively. **C)** SINV-nluc levels, measured by luminescence, upon individual silencing of 461 genes in Aag2 cells. The 2-fold threshold is indicated and putative antiviral and proviral genes are indicated in blue and red, respectively. SINV-nluc infection was performed with MOI = 0.1. Bars are means of three replicates. **D)** Validation of the RNAi screen. In infection with SINV (MOI = 0.1), candidate genes were silenced in Aag2 cells using two independent sets of dsRNA and virus replication was measured with a luminescence assay. **E)** Quantification of SINV RNA levels by RT-qPCR after silencing of the indicated genes in Aag2 cells using the first set of dsRNA. **F)** Infectious DENV-2 titers in the supernatant of Aag2 cells upon *AAEL004419/Dhx15, AAEL008728, AAEL004859* and *AAEL001216* silencing. DENV-2 infection MOI 0.1. **G)** Quantification of CHIKV RNA levels by RT-qPCR after silencing of *AAEL004419/Dhx15, AAEL008728* and *AAEL004859*. CHIKV infection in Aag2 C3PC12 cells was performed with MOI = 0.1. In panels (***D-G***), bars and whiskers represent the mean +/- SD of three independent biologicals replicates. In (***F***) and (***G***), statistical significance was determined using One-Way ANOVA with Holm-Sidak correction (* *p* < 0.05, ** *p* < 0.005, *** *p* < 0.0005).

Here, we focused on the genes that enhanced virus replication upon knockdown, as those are putative players in antiviral defense. Importantly, knockdown of these genes did not, or only mildly, affect cell viability (Fig. S1A). To validate the antiviral activity using an independent readout, we assessed the effect of gene knockdown on SINV replication at the RNA level. Efficient silencing could be verified for most genes (Fig. S1B) and, analogous to our findings measuring luciferase, resulted in an increase of viral RNA levels (Fig. 1E), underscoring the robustness of our screening approach.

Amongst the hits of our RNAi screen, we identified five predicted DEAD-box RNA helicases (AAEL001216, AAEL004419, AAEL004859, AAEL006794, and AAEL008728) amongst which the known antiviral RNAi factor Dicer 2 (AAEL006794) and four RNA helicases that had not previously been associated with antiviral activity in mosquitoes. Given the importance of this class of RBPs in modulating immune signaling, we further focused our analysis on the uncharacterized RNA helicases. First, we aimed to establish the antiviral activity of these DEAD-box helicases against other arboviruses. Silencing of *AAEL004419, AAEL008728* and *AAEL004859*, but not *AAEL001216* resulted in a profound increase of dengue virus titers, to similar levels as silencing of Ago2 (Fig. 1F). Similarly, knockdown of *AAEL004419, AAEL008728* and *AAEL004859* in Aag2-C3PC12 cells, enhanced RNA replication of CHIKV by > 2-fold (Fig. 1G), suggesting a broad antiviral activity of these helicases. C3PC12 cells are an Aag2 cell sub-clone that was cleared from persistently infecting viruses (50). Importantly, silencing of the identified RNA helicases in these cells resulted in increased Sindbis virus levels as observed in the initial knockdown screen, which had been performed in the parental Aag2 cell line (Fig. S1C and D).

### Characterization of broadly antiviral RNA DEAD-box helicases

AAEL004419, AAEL008728 and AAEL004859 are canonical DEAD-box helicases containing DEAD-like helicase superfamily (DEXDc) and helicase superfamily C-terminal (HELICc) domains. In addition, AAEL004419 and AAEL004859 contain a C-terminal helicase associated (HA2) domain and AAEL004859 contains two double stranded RNA binding motifs (DSRM) (Fig. 2A). Alignment of *Ae. aegypti* and *Drosophila* DEAD-box helicase domains, identified Dhx15, CG9143, and maleless (mle) as the closest orthologs of AAEL004419, AAEL008728, and AAEL004859, respectively (Fig. S2A). In particular, AAEL004419 is highly conserved with about 90% amino acid identity across all functional domains (Fig. S2B). Because of the close one-to-one orthology, we will refer to AAEL004419 as Dhx15.

**Figure 2:**
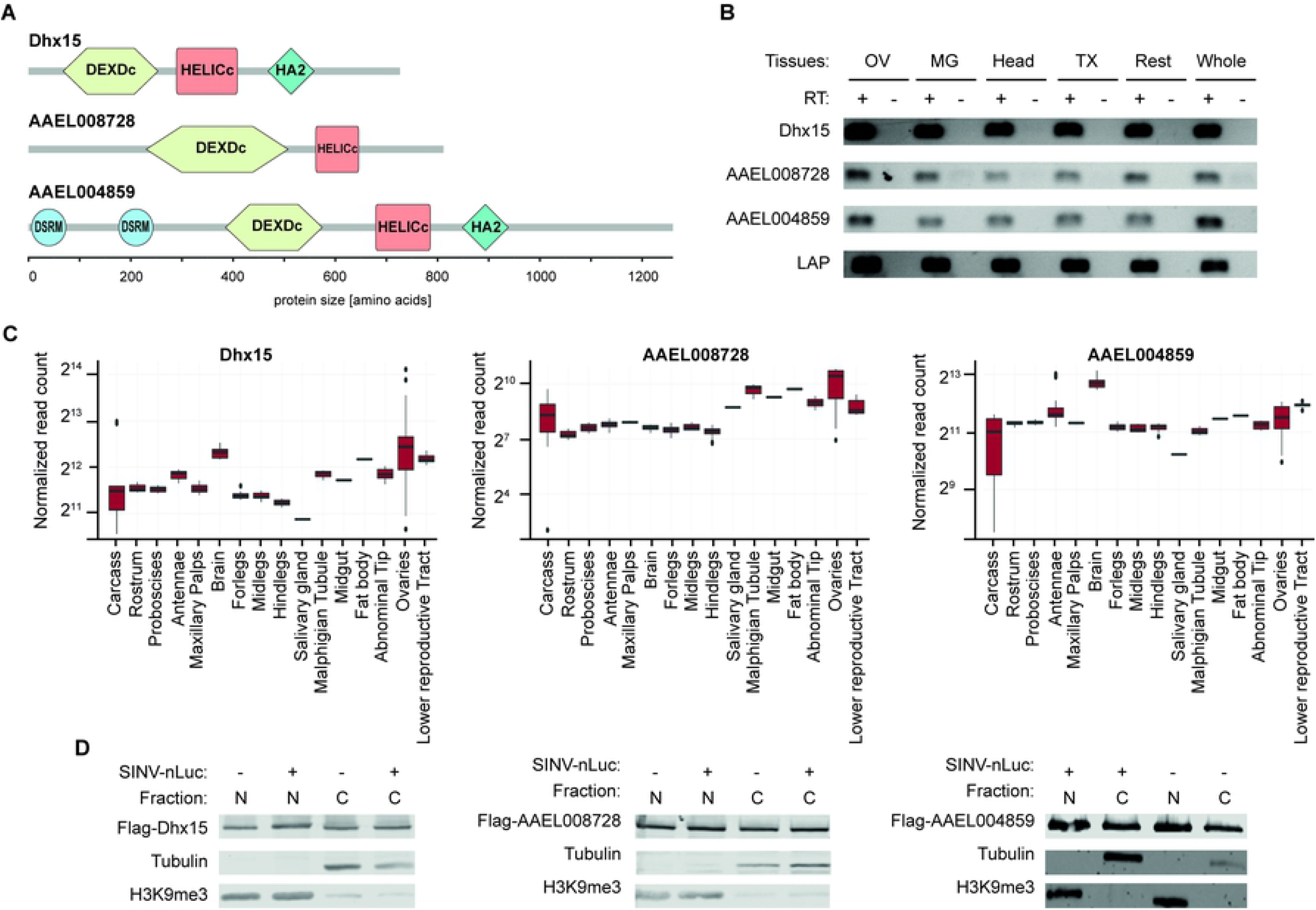
Characterization of AAEL004419/Dhx15, AAEL008728 and AAEL004859. **A)** Schematic representation of the domain structure of the RNA helicases AAEL004419/Dhx15, AAEL008728 and AAEL004859 predicted with SMART. DEDXc: DEAD-like helicase superfamily domain, HELICc: helicase superfamily C-terminal domain, HA2: C-terminal helicase associated domain, DSRM: Double stranded RNA binding motif. **B)** Expression of AAEL004419/Dhx15, AAEL008728, AAEL004859 and the house-keeping gene Lysosomal Aspartic protease (LAP) assessed by RT-PCR on ovaries (OV), midgut (MG), head, thorax (TX), rest of the body dissected from female *Ae. aegypti* mosquitoes as well as in entire mosquitoes. PCR amplification on samples without reverse transcriptase (RT−) served as negative control. **C)** Expression of AAEL004419/Dhx15, AAEL008728 and AAEL004859 in mosquito tissues in published RNA-seq datasets. **D)** Cellular localization of the proteins of interest in noninfected (-) and SINV infected (+) Aag2 C3PC12 cells. SINV-nLuc infection was performed at MOI = 0.1. Cell fractionation assay followed by western blot show the expression of AAEL004419/Dhx15, AAEL008728 and AAEL004859 in the nucleus (N) and in the cytoplasm (C).

To further characterize the three DEAD-box helicases, we investigated their expression pattern both at the tissue level in adult mosquitoes and on and sub-cellular level in Aag2 cells. In dissected female *Ae. aegypti* mosquitoes, we found Dhx15, AAEL008728, and AAEL004859 to be ubiquitously expressed across all somatic and germline tissues analyzed (Fig. 2B), which is in line with published RNA expression data (Fig. 2C). To assess the subcellular localization of Dhx15, AAEL008728, and AAEL004859, we expressed Flag-tagged proteins in *Ae. aegypti* Aag2 cells and performed nuclear versus cytoplasmatic fractionation. Efficient separation of the cytoplasmic and nuclear fractions was confirmed by the segregation of tubulin and Histone3-Lysine 9 tri-methylation (H3K9me3) markers, respectively. We identified all three RNA helicases to be ubiquitously expressed in the nuclear and in the cytoplasmic fractions both in uninfected and SINV infected Aag2 cells, indicating that subcellular localization was not altered as a response to virus infection (Fig. 2D).

### Silencing of *Dhx15* results in an altered transcriptional response regulating glycolysis

In vertebrates, orthologues of *Dhx15, AAEL008728* and *AAEL004859* have been proposed to regulate transcriptional responses to virus infection by modulating signal transduction of core immune pathways such as MAPK (mitogen-activated protein kinase) and NFκB signaling (51–54). We therefore decided to investigate transcriptional regulation mediated by the highly conserved RNA helicase *Dhx15*. After sequential knockdown of *Dhx15* in C3PC12 cells, we performed RNA-sequencing and gene expression analysis (Fig. 3A). Genes were considered differentially expressed (DE) when their expression levels were up or down-regulated by at least 2-fold and the adjusted *p*-value was *p* < 0.05. Using these parameters, we identified 528 genes upregulated and 328 genes downregulated upon *Dhx15* knockdown (Fig. 3B). For the up-regulated genes, GO terms related to DNA replication were the most strongly enriched; for the downregulated genes, GO terms related to sugar metabolism, most prominently *glycolytic process*, were the most strongly enriched (Fig. 3C). These results indicate that *Dhx15* directly or indirectly controls a transcriptional response in Aag2 cells.

**Figure 3:**
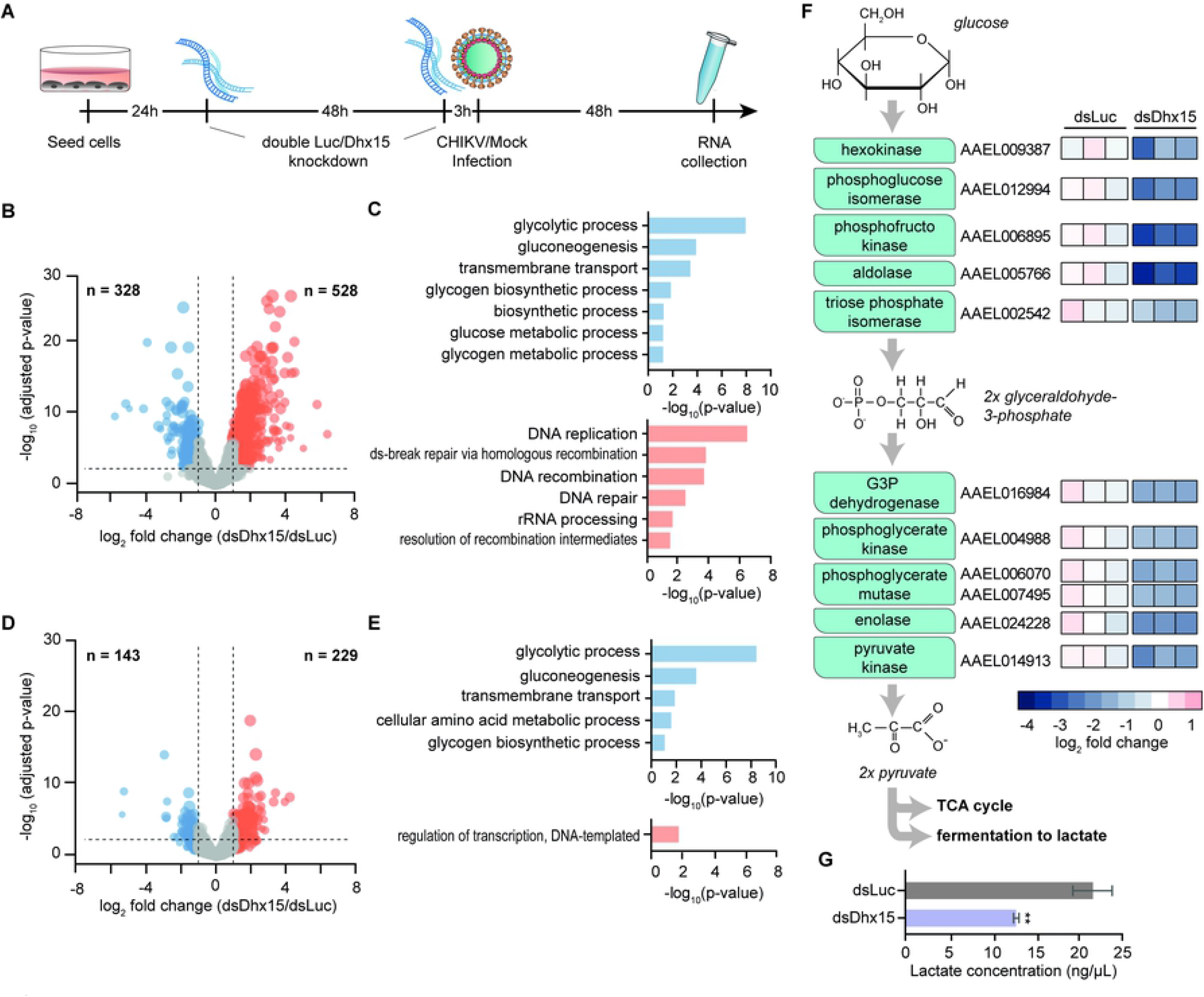
*Dhx15* regulates a transcriptional response that controls glycolysis. **A)** Set-up of RNA-seq analysis to assess the transcriptomic response to *Dhx15* silencing. 24 hours and 48 hours after Aag2 C3PC12 cells were seeded, a sequential knockdown of *Dhx15* (dsDhx15) or a non-targeting Firefly luciferase (dsLuc) control was performed. CHIKV (MOI = 5) or mock infection was performed 3 hours after the second knockdown and samples were collected 48 hours later. **B)** Volcano plot of differential expression of *Dhx15* silencing depicting comparison between downregulated genes (blue) and upregulated genes (red). The X-axis denotes log2 fold change values; the Y-axis shows −log10 (*P*-value). **C)** GO terms of differentially expressed genes upon *Dhx15* silencing. Upper panel (blue), GO analysis of downregulated genes. Lower panel (red), GO annotation of upregulated genes. **D)** Volcano plot of *Dhx15* silencing in the context of CHIKV infection, showing downregulated genes and upregulated genes in blue and red, respectively. The X-axis denotes log2 fold change values; the Y-axis shows −log10 (*P*-value). **E)** GO annotation of differentially expressed genes upon *Dhx15* silencing and CHIKV infection. Upper panel (blue), downregulated genes. Lower panel (red), upregulated genes. **F)** Schematic representation of the players involved in the glycolysis pathway (left) and log_2_ fold change of these genes upon *Dhx15* or Firefly luciferase silencing (right). **G)** Relative lactate concentration upon *Dhx15* or Firefly luciferase silencing in Aag2 C3PC12 cells. Bars and whiskers represent the mean +/- SD of three independent biologicals replicates. Statistical significance was determined using unpaired two tailed t-test (** *p* < 0.005).

We next assessed the effect of *Dhx15* knockdown on gene expression in the context of virus infection. Aag2 cells were infected with CHIKV shortly after the second knockdown, and RNA samples were taken 48 hours post infection. Efficient CHIKV replication and *Dhx15* knockdown were verified in these samples (Fig. S3A and B) and analysis of RNA deep-sequencing data identified 229 genes and 143 genes to be significantly up or downregulated, respectively (Fig. 3D). The majority of these (194 out of 229 upregulated genes and 89 out of 143 downregulated genes) overlapped with the differentially expressed genes in uninfected samples, defining a set of genes with robust *Dhx15*-dependent differential expression, regardless of virus infection (Fig. S3C). Interestingly, while for the up-regulated genes, *DNA templated regulation of transcription* was the only enriched GO term, for the downregulated genes, GO terms were highly concordant between uninfected and infected conditions with *glycolytic process* being the most strongly enriched (Fig. 3E). We therefore specifically analyzed the expression of genes that are part of the glycolysis pathway, and indeed found that the entire set of glycolytic core enzymes was downregulated upon *Dhx15* knockdown, in particular those that are involved in the metabolic conversion of glucose to glyceraldohyde-3-phosphate (Fig. 3F and S3D).

Amongst the most strongly downregulated genes is the gene encoding phosphofructokinase, the enzyme that performs the rate-limiting step of the glycolysis pathway. We therefore functionally assessed the effect of *Dhx15* knockdown on the glycolytic rate by measuring the concentration of lactate, a fermentation product of the glycolysis pathway as a proxy for activity (55–57). To benchmark our assay, we treated Aag2 cells with 2-deoxy-D-glucose (2-DG), which is converted by hexokinase into 2-deoxy-D-glucose phosphate, a competitive inhibitor of phosphoglucose isomerase at the second step of glycolysis (58). As a control, we treated Hela cells, for which 2-DG treatment is known to reduce lactate concentration (59). As expected, treatment with 2-DG resulted in an almost 30% decline of lactate levels in Hela cells (Fig. S3E). In contrast, in Aag2 cells, baseline lactate levels were lower and treatment with 2-DG only had a minor effect on lactate concentration (Fig. S3F). We hypothesized that this may be explained by the composition of the L-15 culture medium, which contains galactose instead of glucose and additional high levels of pyruvate. Galactose can enter glycolysis but at lower efficiency than glucose and high levels of pyruvate favor energy production by directly entering into the tricarboxylic acid cycle, which likely reduces the glycolytic activity to form lactate. To sensitize our lactate assay, we therefore cultured Aag2 cells in Schneider’s medium, which is supplemented with 11.11 mM glucose and does not contain pyruvate. In these culture conditions, baseline lactate levels were elevated, and 2-DG treatment resulted in significantly lower lactate concentrations, indicating that we were able to measure alterations in glycolytic activity in Aag2 cells (Fig. S3F). We next assessed lactate levels upon *Dhx15* silencing. Strikingly, we observed a profound decrease of lactate concentration in cell homogenates, even exceeding the effect of 2-DG treatment, indicating that the reduced expression of glycolysis genes upon *Dhx15* knockdown results in a functional reduction of glycolytic activity in Aag2 cells (Fig. 3G). The decrease in lactate concentration cannot be explained by a reduced cell number, which remained stable or was slightly elevated upon *Dhx15* silencing (Fig. S3G). Altogether, our results suggest that *Dhx15* knockdown effectively downregulates mRNA expression of core glycolytic enzymes resulting in functional reduction of glycolysis rate.

### Transcriptional control of glycolytic genes is specific to Dhx15

We next aimed to investigate whether, besides Dhx15, the other identified antiviral DEAD-box helicases contributed to transcriptional downregulation of glycolytic genes. This hypothesis was sparked by a protein-protein interaction map that we generated for all 15 antiviral hits picked up in our screen using the STRING algorithm. In this analysis, all identified DEAD-box-helicases were predicted to interact in a protein complex (Fig. 4A). To confirm a direct protein-protein interaction of AAEL008728 with Dhx15 and AAEL004859 experimentally, we performed co-immunoprecipitations (Co-IP) in Aag2 cells. Confirming the predicted protein interaction network, Flag-tagged AAEL008728 was efficiently co-precipitated both by GFP-tagged AAEL004859 and Dhx15 (Fig. 4B). Since the three DEAD-box helicases are predicted to have RNA binding activity, it was plausible that their interaction was mediated indirectly through binding to the same RNA molecules. To explore this possibility, we performed Co-IP in the presence of RNase A to disrupt RNA-bridged protein-protein interactions. Dhx15 and AAEL008728 binding was resistant to RNase A treatment (Fig. 4C), indicating an RNA-independent interaction between these helicases.

**Figure 4:**
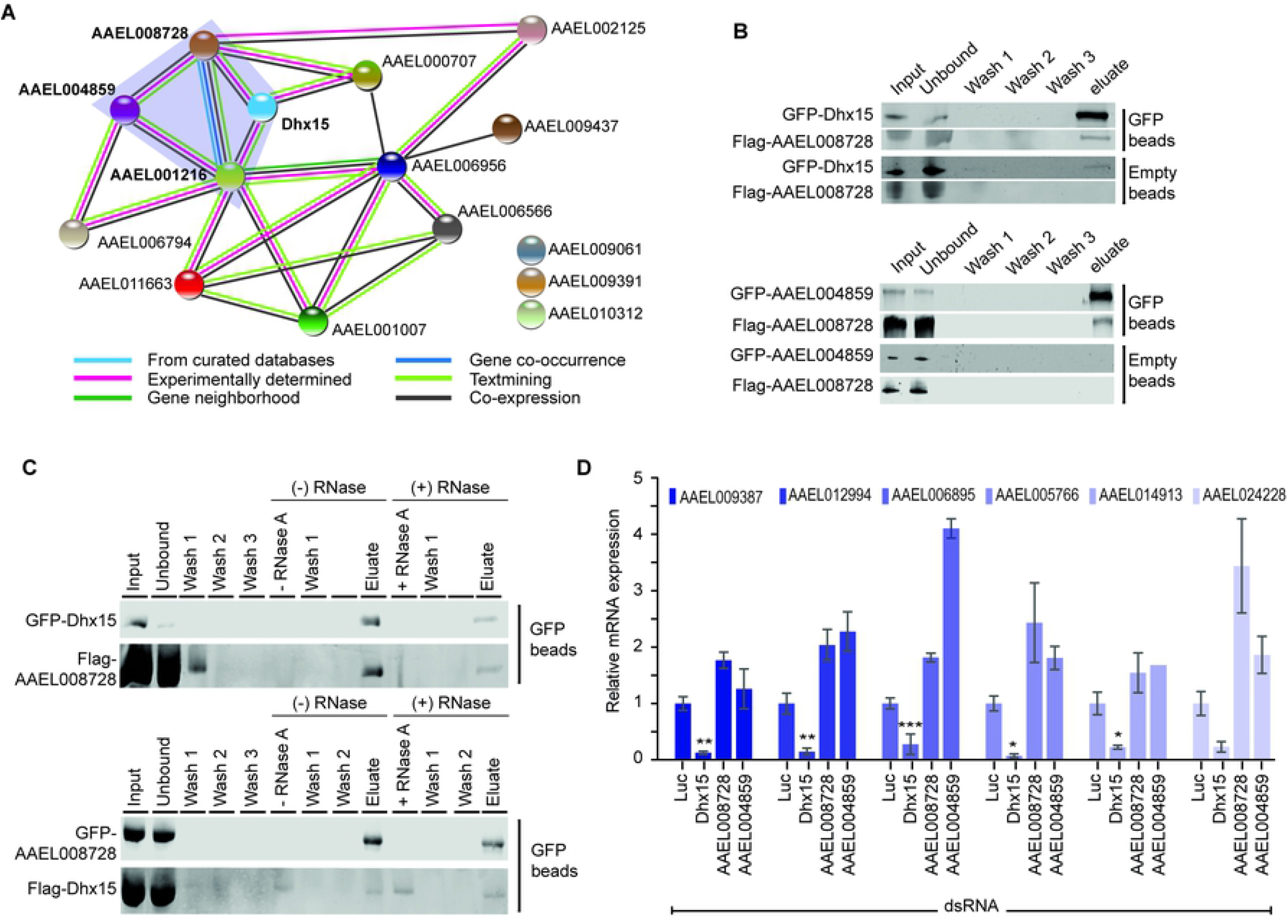
*Dhx15* silencing, but not other RBPs downregulate glycolytic genes. **A)** Protein-protein interactions predicted for fifteen antiviral RBPs using STRING. The color code of the lines connecting the different RBPs, represents the prediction for the protein-protein association. A network of uncharacterized DEAD-box RNA helicases is highlighted with a purple background. **B)** Western blot analysis of protein lysates from Aag2 cells transfected with GFP-Dhx15 and Flag-AAEL008728 (top panel) as well as GFP-AAEL004859 and Flag-AAEL008728 (bottom panel). Samples before (input) and after GFP-IP or control IP with empty beads were analyzed for co-purification of GFP- and Flag-tagged transgenes. Samples were probed with antibodies against GFP and Flag. **C)** Co-IPs of GFP-Dhx15 and Flag-AAEL008728 (top panel) and GFP-AAEL008728 and Flag-Dhx15 (bottom panel) from Aag2 C3PC12 cell lysate with (+) and without (-) subsequent on-bead RNase A treatment. RNase A was added to the sample after the 3 initial wash steps post IP and samples taken directly after incubation are denoted as – RNase A and + RNase A in vertical writing, respectively. Samples were probed with antibodies against GFP and Flag. **D)** Quantification of glycolytic genes by RT-qPCR after silencing of *Dhx15, AAEL008728* and *AAEL004859* in Aag2 C3PC12 cells. Bars and whiskers represent the mean +/- SD of three independent biologicals replicates. Statistical significance was determined using One-Way ANOVA with Holm Sidak correction (* *p* < 0.05, ** *p* < 0.005, *** *p* < 0.0005).

Having confirmed a direct interaction of the identified DEAD-box helicases, we next assessed if knockdown of AAEL008728 and AAEL004859 caused a similar transcriptional response as Dhx15. Quantification of glycolytic genes and an additional selection of differentially regulated genes from the RNA-seq data confirmed the reduced expression upon silencing of *Dhx15*. However, knockdown of *AAEL008728* and *AAEL004859* did not reduce expression of these genes (Fig. 4D and S4A), indicating that the transcriptional control of glycolysis genes was not mediated by the protein complex of the three helicases identified (Fig. 4A) but rather by a function specific to Dhx15.

### CHIKV infection down-regulates glycolysis genes, akin to *Dhx15* knockdown

In response to virus infections, the activity of metabolic pathways is often changed reflecting the altered energy and biomolecule demand in infected cells (33). Therefore, we wanted to assess the general transcriptional response of Aag2 cells to CHIKV infection. For this aim, we re-analyzed our RNA-seq datasets comparing gene expression of uninfected and CHIKV infected Aag2 cells treated with non-targeting control dsRNA. This analysis allowed us to identify genes that are differentially regulated in response to CHIKV infection in Aag2 cells irrespective of *Dhx15* knockdown. In general, the transcriptional response to CHIKV in Aag2 cells was modest with only 8 genes upregulated and 51 genes downregulated (Fig. 5A). Strikingly, amongst the 51 downregulated genes, a significant number of genes (n=22; Pearson Chi-square < 0.001) were also consistently decreased by *Dhx15* knockdown (Fig. 5B), suggesting that CHIKV infection and *Dhx15* knockdown results in a partially overlapping transcriptional response. CHIKV induced gene repression was not mediated by downregulation of *Dhx15*, as expression of this RNA helicase was not altered in infected cells (Fig. 5C). Strikingly, GO analysis identified *glycolytic process* as the only enriched term (Fig. 5A). Three core enzymes of the glycolysis, aldolase, hexokinase, and the rate-limiting phosphofructokinase were significantly downregulated (Fig. 5A and D). More general, all eleven glycolytic genes are expressed at a lower level in infected cells, albeit not always reaching our thresholds for minimal fold changes or statistical significance (Fig. S5A). Particularly, the expression of enzymes involved in the first metabolic steps of glycolysis are reduced, mimicking the effect of *Dhx15* knockdown (Fig. 3F and S5A). These data suggest that silencing of *Dhx15* is involved in regulating a glyco-metabolic response that establishes a cellular environment that favors CHIKV replication, likely through alterations in metabolic rates or synthesis of precursors of biomolecules.

**Figure 5:**
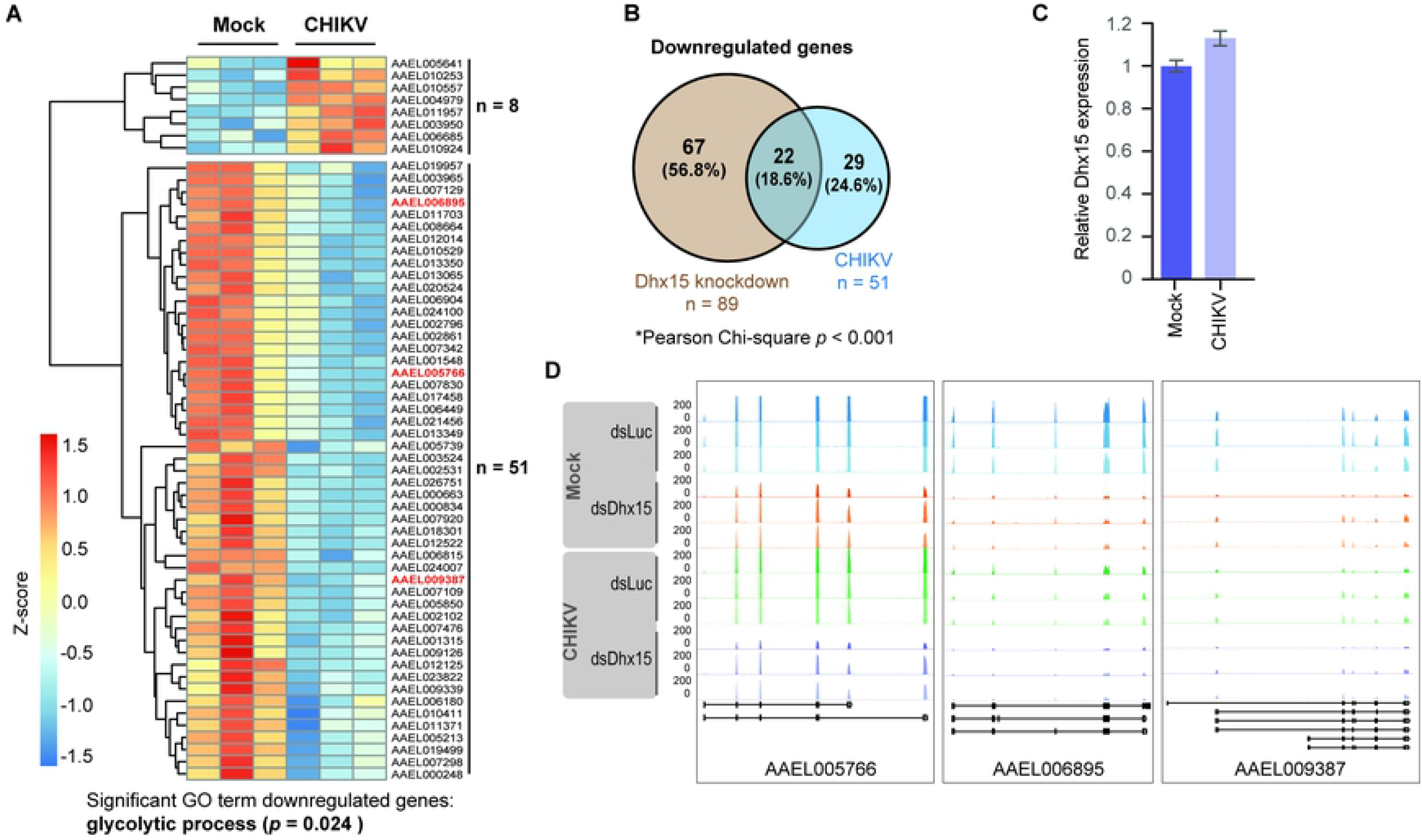
Glycolytic genes are downregulated upon CHIKV infection. **A)** Heatmap of differentially expressed genes upon CHIKV infection MOI 5 (fold change ≥ 2; *p*-value < 0.05). Z-score was calculated based on log10 fold changes of each gene to indicate the level of expression. **B)** Overlap of genes downregulated by *Dhx15* silencing and CHIKV infection as identified by RNA-seq. Statistical significance was determined using Pearson Chi-square test (*p* < 0.001). **C)** Relative expression of *Dhx15* in uninfected and CHIKV infected cells, extracted from RNA-seq data. **D)** RNA-seq tracks for AAEL05766 (aldolase) AAEL006895 (phosphofructokinase) and AAEL009387 (hexokinase) from the indicated conditions.

## Discussion

*Aedes aegypti* mosquitoes are important biological vectors for major arboviruses that impose a growing threat to human health (68), asking for a better understanding of the mechanisms that control virus growth in mosquitoes. Similar to other insect species, antiviral immunity in mosquitoes is mediated through small RNA-mediated silencing of viral RNA as well as transcriptional responses to virus infections (17). While the antiviral mechanisms underlying small RNA pathways, in particular RNAi, are relatively well established, mechanistical insights into how transcriptional responses govern antiviral immunity are limited (15–17), and additional, yet unknown, pathways that control virus replication in mosquitoes likely exist. In order to identify new players in antiviral defense, we performed a targeted knockdown screen in Aag2 mosquito cells, a cell line of embryonic origin that is frequently used to molecularly dissect antiviral immune pathways (21, 60, 61). We focused this functional screen on RBPs, a protein family with pleiotropic functions in regulating immune responses across all domains of life (23–27). Indeed, using a robust screening strategy including two confirmation rounds with independent dsRNA sequences, we identified several proteins with antiviral properties that, when silenced, resulted in increased virus replication. Amongst the hits were the well-established antiviral RNAi factor Dicer 2 (AAEL006794) and proteins that act in transcriptional pausing (Spt4: AAEL006566 and Spt6: AAEL006956), a process that had previously been reported to have antiviral activity in flies and Aag2 mosquito cells (62).

From the identified hits, we initially focused on DEAD-box helicases and in particular the role of Dhx15. Dhx15 exhibits broad antiviral activity against SINV and CHIKV, two viruses of the alphavirus genus and DENV, a flavivirus. We show that *Dhx15* controls a transcriptional response that decreases glycolytic activity in mosquito cells. Intriguingly, CHIKV infections results in a similar reduction of genes involved in the glycolysis pathway. Although further mechanistic experiments are needed, our data suggest that the enhanced virus replication of CHIKV upon *Dhx15* knockdown may be explained by establishment of a metabolic environment that favors CHIKV replication. Thus, while our RNAi screen was initially intended to discover new immune factors that directly interfere with virus replication, we eventually identified a protein that indirectly represses virus growth potentially by controlling the metabolic state of the cell.

DEAD box helicases have a large array of functions in general RNA metabolism as well as controlling antiviral immunity (63–65). Prime examples are the DEAD-box helicases Dicer-2 and the RIG-I like RNA helicases, which are essential for sensing viral RNA in invertebrates and vertebrates, respectively (24, 29). In addition, DEAD box helicases modulate immune signaling via direct interaction with core signaling intermediates in the cytoplasm or by regulating transcription in the nucleus as coactivators or co-suppressors of transcription factors (66, 67). As such, several DEAD box helicases in mammals (i.e.: DDX1, DDX3, DHX9, DHX15, DDX21, DDX24 DHX33 and DHX36) exert broadly antiviral effects against a variety of RNA and DNA viruses (51, 53, 54, 67–71). In line with this, we identified three DEAD box helicases, Dhx15, AAEL004859 and AAEL008728, to also have a broad antiviral phenotype against SINV, DENV and CHIKV infections.

We pursued an in-depth characterization of Dhx15, a highly conserved DEAD-box RNA helicase that was previously characterized as a part of the U2 spliceosome in vertebrates and invertebrates (72). Interestingly, in human cells Dhx15 acts as a co-receptor for Rig-I like receptors (RLR) and is required for antiviral RLR signaling (71). Furthermore, Dhx15 activates MAPK and NFκB signaling during antiviral responses triggered by poly I:C and the two RNA viruses encephalomyocarditis and Sendai virus (51). We, therefore, speculated that *Dhx15* is involved in regulating a transcriptional response in mosquito cells, as well. Indeed, *Dhx15* silencing caused hundreds of genes to be differentially expressed, both in uninfected as well as CHIKV-infected cells. However, we did not observe canonical immune target genes to be differentially regulated upon *Dhx15* knockdown. Instead, we observed that all genes that encode enzymes involved in the core glycolysis pathway were consistently downregulated, both in uninfected and CHIKV infected cells. This downregulation resulted in reduced lactate production, suggesting impairment of glycolytic activity in *Dhx15*-depleted mosquito cells. Also in mice, *Dhx15* has previously been linked to energy metabolism but in contrast to our deep-sequencing data, various glycolytic genes were transcriptionally upregulated upon Dhx15 knockdown in mouse endothelial cells (73). It is currently unclear what explains this discrepancy, but it is likely that overall differences in the metabolic state and integration of other regulatory mechanisms within the different experimental systems account for various metabolic outcomes. In this context it is important to note that it is currently unclear via which signaling cascade Dhx15 regulates glycolytic gene expression. In human cancer cells, inhibition of NFκB signaling reduced glycolysis via transcriptional regulation of hexokinase 2, the first enzyme of glycolysis (74). Interestingly, human Dhx15 has been previously shown to activate NFκB signaling, providing a possible link between Dhx15 expression and alterations in glycolytic rates (51). In mosquitoes, two NFκB-like signaling pathways, Toll and IMD, exist (17), and it will be interesting to investigate whether these are involved in the Dhx15 mediated gene expression.

Although the exact mechanism of antiviral activity of Dhx15 remains to be established, we propose that knockdown of *Dhx15* establishes a metabolic environment, in particular through the repression of glycolytic genes, that favors CHIKV infection. Glycolysis can be a relevant source of ATP and supports cell growth by providing intermediates for several biosynthesis pathways. For example, the product of the first enzymatic step of glycolysis, glucose-6-phosphate enters the pentose phosphate pathway (PPP), which is responsible for generating pentoses (five-carbon sugars) as well as other RNA and DNA precursors (31). Changes in glycolytic rate are known to occur widely during virus infection presumably as a consequence of a higher energy and/or nucleotide demand enforced by virus replication (32, 33, 75–77). On the other hand, an increased energy metabolism has been shown to activate antiviral defense and glycolytic enhancement is dispensable or even avoided due to triggering of immune responses of the host (31, 78, 79). Therefore, the activity of metabolic pathways is likely regulated at multiple levels, potentially explaining why the outcome of changing metabolic rates appears to be highly specific for distinct virus-host combinations (32, 77). For example, for alphaviruses, increased glycolytic activity has been proposed to support the elevated demand of cellular energy and biomolecules required during Semliki Forest virus, Mayaro virus (MAYV), and SINV replication (75, 79–81). For CHIKV, however, the effect of the virus infection on the metabolic pathways is dependent on the experimental system (32). On the one hand, CHIKV infection in human cells and in a mouse model incremented cellular metabolism by upregulation of PKM2 and PDHA1, an isoenzyme of pyruvate kinase and a component of the pyruvate dehydrogenase enzyme complex, respectively (82, 83). On the other hand, CHIKV infection lead to downregulation of glycolytic enzymes in a human hepatic cell line (82), similar to what we have observed in *Ae. aegypti* cells.

While in vertebrates, CHIKV virus infections cause a dramatic change in gene expression profiles, largely as a consequence of immune gene induction upon stimulation of interferon signaling (84), we found that CHIKV infection in Aag2 cells only resulted in differential expression of a few dozen genes, the vast majority of which was downregulated. It is currently not clear what causes this curiously weak transcriptional response; two non-mutually exclusive hypotheses are a generally more delicate transcriptional immune signaling in Aag2 cells or active suppression of transcriptional responses by CHIKV and possibly other arboviruses. Despite the modest transcriptional response to CHIKV infection, there was a remarkable overlap of genes that were downregulated upon *Dhx15* silencing and CHIKV infection. It is tempting to speculate that *Dhx15* knockdown creates a metabolic environment that mimics CHIKV infection thereby allowing for enhanced virus replication. Altogether, our results uncover an intriguing interaction between transcriptional regulation mediated by a host DEAD-box RNA helicase, alterations in metabolic activities, and antiviral activity in mosquito cells.

## Acknowledgements

The authors would like to thank Rebecca Halbach for analyzing expression of DEAD-box helicases in published sequencing data. Thanks also to Ronald van Rij for critical reading of the manuscript. The CHIKV Leiden synthetic LS3 construct was kindly provided by Dr. Martijn J van Hemert at Leiden University Medical Center. Thanks to BEI resources established by the National Institute of Allergy and Infectious Diseases for providing *Aedes aegypti* Liverpool mosquitoes. This work was financially supported by a Veni grant (ID: VI.Veni.202.035) from the Dutch Research Council (Nederlandse Organisatie voor Wetenschappelijk Onderzoek; NWO) to PM.

## Authors contribution

SRM and PM conceptualized the project. SRM and JQ performed the experiments. JQ performed the bioinformatics analyses. SRM, JQ and PM analyzed and interpreted the data. WK analyzed data of metabolic experiments. SRM and PM wrote the manuscript, all authors read and edited the paper. PM acquired funding.

## Figures

**Figure S1: RBP candidate genes that control arboviruses replication in mosquito cells A)** Viability of Aag2 cells was measured using CellTiter-Glo assay after silencing of 15 candidate genes (see Fig. 1D and E) using the first set of double-stranded RNA. Bars and whiskers represent the mean +/- SD of three independent biologicals replicates. **B)** Knockdown efficiency of 15 candidate genes (from experiment shown in Fig. 1E) was assessed by RT-qPCR. Bars and whiskers represent the mean +/- SD of three independent biologicals replicates. Statistical significance was determined using unpaired two tailed t-test (* *p* < 0.05, ** *p* < 0.005, *** *p* < 0.0005). **C)** Levels of SINV were quantified using RT-qPCR after silencing of *Dhx15, AAEL008728* and *AAEL004859* in Aag2 C3PC12 cells. SINV infection with MOI 0.1. Bars and whiskers represent the mean +/- SD of three independent biologicals replicates. Statistical significance was determined using One-Way ANOVA with Holm Sidak correction (*** *p* < 0.0005). **D)** Knockdown efficiency of genes from panel (***C***) were assessed by RT-qPCR. Bars and whiskers represent the mean +/- SD of three independent biologicals replicates. Statistical significance was determined using unpaired two tailed t-test (* *p* < 0.05).

**Figure S2: AAEL004419 is the direct orthologue of Dhx15. A)** Unrooted approximately-maximum likelihood tree of *Drosophila* (purple) and *Ae. aegypti* (brown) RNA-helicases with branch lengths estimated using the CAT approximation described in (49). **B)** Multiple sequence alignment of *Drosophila* Dhx15 and *Ae. Aegypti* AAEL004419. The functional domains (see Fig. 2A) are highlighted with colored boxes.

**Figure S3: RNA-seq analysis identifies *Dhx15* as regulator of glycolysis. A-B)** Levels of CHIKV (***A***) and knockdown efficiency of *Dhx15* (***B***) in samples used for deep-sequencing were assessed by RT-qPCR. CHIKV infection was performed with MOI = 5. Bars and whiskers represent the mean +/- SD of three independent biologicals replicates. Statistical significance was determined using unpaired two tailed t-test (** *p* < 0.005, *** *p* < 0.0005). **C)** Number of overlapping genes downregulated (left panel) and upregulated (right panel) upon *Dhx15* silencing in uninfected and CHIKV infected cells. **D)** Schematic representation of the players involved in the glycolysis pathway (left) and log_2_ fold change of these genes upon *Dhx15* or Firefly luciferase silencing (right) in CHIKV infected cells. **E-F)** Relative lactate concentration upon 2-deoxy-D-glucose (2-DG) treatment in Hela (***E***) and Aag2 (***F***) cells. **G)** Number of Aag2 cells after sequential *Dhx15* or Firefly luciferase knockdown. Bars and whiskers represent the mean +/- SD of three independent biologicals replicates. Statistical significance was determined using unpaired two tailed t-test (* *p < 0*.*05*, *** *p* < 0.0005).

**Figure S4: Differentially regulated genes derived from RNA-seq data are specifically dependent on *Dhx15* silencing. A)** Quantification of top five most differentially regulated genes obtained from the RNA-seq list of 22 genes with shared downregulation between CHIKV infection and *Dhx15* knockdown (Fig. 5B). RNA levels were measured by RT-qPCR after individual silencing of *Dhx15, AAEL008728* and *AAEL004859* in Aag2 C3PC12 cells. Bars and whiskers represent the mean +/- SD of three independent biologicals replicates. Statistical significance was determined using One-Way ANOVA with Holm Sidak correction (* *p* < 0.05, *** *p* < 0.0005).

**Figure S5: CHIKV infection causes reduction of glycolytic genes. A)** Relative expression of genes from the glycolysis pathway in CHIKV infected cells (MOI = 5) compared to uninfected cells in control (dsLuc) knockdown conditions. Expression values were extracted from the RNA-seq data and normalized to uninfected cells. Bars and whiskers represent the mean +/- SD of three independent biologicals replicates. Statistics from the DESeq2 analysis are shown (* *P adj < 0*.*05*, (** *P adj* < 0.005, *** *P adj* < 0.0005).

**Table S1: Oligonucleotides used in this study**.

**Table S2: Raw data from target knockdown screen and confirmation rounds**. Genes that have been selected for an RNAi screen and, if applicable, updated gene identifiers in the recent version of VectorBase (version 57, accessed April 2022).

**Table S3: Differentially expressed genes in different comparisons**. List 1: *Dhx15* knockdown vs. control knockdown in uninfected cells. List 2: *Dhx15* knockdown vs. control knockdown in CHIKV cells. List 3: CHIKV infected vs. uninfected cells in control knockdown conditions.

**Table S4: Source data file**.

## Notes

### Competing Interest Statement

The authors have declared no competing interest.

